# The energetics of social signaling during roost location in Spix’s disc-winged bats

**DOI:** 10.1101/2020.09.24.312496

**Authors:** Gloriana Chaverri, Paula Iturralde-Pólit, Natalia Ivone Sandoval-Herrera, Adarli Romero-Vásquez, Silvia Cháves-Ramírez, Maria Sagot

**Affiliations:** Sede del Sur, Universidad de Costa Rica, Golfito, CRI; Smithsonian Tropical Research Institute, Balboa, Ancón, PAN; Department of Ecology and Evolutionary Biology, University of Toronto, Ontario, CAN; Escuela de Biología, Universidad de Costa Rica, San Pedro, CRI; Escuela de Biología, Universidad Nacional, Heredia, CRI; Department of Biological Sciences, State University of New York at Oswego, New York, USA

**Author notes:** These authors contributed equally (Paula Iturralde-Pólit, Natalia Ivone Sandoval-Herrera).

**Keywords:** allocation model, bats, energetic expenditure, resting metabolic rate, social calls

## Abstract

Long-term social aggregations are maintained by multiple mechanisms, including the use of acoustic signals, which may nonetheless entail significant energetic costs. To date, however, no studies have gauged whether there are significant energetic costs to social call production in bats, which heavily rely on acoustic communication for a diversity of social tasks. We measure energetic expenditure during acoustic signaling in Spix’s disc-winged bats (*Thyroptera tricolor*), a species that commonly uses contact calls to locate the ephemeral furled leaves that they use for roosting. To determine the cost of sound production, we measured oxygen consumption using intermittent-flow respirometry methods, with and without social signaling. Our results show that the emission of contact calls significantly increases oxygen consumption; vocal individuals spent, on average, 12.42 kJ more during social signaling trials than they spent during silent trials. Furthermore, production of contact calls during longer periods increased oxygen consumption for males but not for females. We also found that as resting metabolic rates increased in males, there was a decreasing probability that they would emit response calls. These results provide support to the “allocation model”, which predicts that only individuals with lower self-maintenance costs can afford to spend energy in additional activities. Our results provide a step forward in our understanding of how physiology modulates behavior, specifically how the costs of call production and resting metabolic rates may explain the differences in vocal behavior among individuals.

**Summary Statement:** Spix’s disc-winged bats constantly produce contact calls while searching for roosts, which we show significantly increases an individual’s metabolic rate.

## Introduction

Many social animals rely on acoustic signals to facilitate social coordination (Kondo and Watanabe 2009; Fichtel and Manser 2010). In bats, for example, social calls are used to locate dependent young, mating partners, prompt and coordinate cooperative interactions, and/or defend and announce the location of resources, including roosts (Chaverri et al. 2018). The latter is of critical importance given that roosts provide refuge from predators and inclement weather, and are the main sites where social interactions, such as lactation, grooming, and mating, occur (Kunz 1982). Thus, the use of social calls during roost finding increases the probability of engaging in beneficial social interactions while reducing the risks of predation; as such, these acoustic signals represent a critical component of social living.

Despite our growing understanding of the benefits of social signaling, particularly in bats, we still do not understand its costs in different contexts. Studies in other taxa suggest that vocalizations that serve a social function increase an individual’s risk of being detected by predators (Magrath et al. 2010) or by potential prey (Deecke et al. 2005), which could reduce foraging efficiency. Moreover, the production of acoustic signals may also carry significant metabolic costs. For example, energy expenditure of vocalizing animals could be up to eight times higher than those of silent ones (Ophir et al. 2010). In bats, echolocation calls produced during flight carry no additional energetic costs beyond those required to power flight (Speakman and Racey 1991; Voigt and Lewanzik 2012), yet may entail significant metabolic costs when produced while roosting, likely due to the contraction of muscles involved in sound emission (Dechmann et al. 2013). However, despite the costs of sound production, the benefits to group coordination and roost-finding efficiency are significant, as just a few calls produced by a single roosting bat are enough to maintain group cohesion and decrease the time needed to locate a new roost site (Sagot et al. 2018).

The costs of call production may potentially explain why social calls are not emitted more frequently, in specific contexts, or by all group members. In moving groups, for example, members may produce social calls only sporadically (Deecke et al. 2005), and individuals may become silent altogether when faced with increased levels of predation risk (Abbey-Lee et al. 2016). The energetic costs of sound production may also explain why only some group members vocalize, as has been observed in bats where lactating females produce significantly fewer calls compared to non-reproductive and pregnant females (Chaverri and Gillam 2015). These intraspecific differences suggest that vocalizations involve higher energetic costs and that non-energetically limited individuals may be able to afford sound production for social communication.

Here, we aim to estimate the energetic cost of social calling in roosting bats to understand patterns of inter-individual differences in vocal behavior. We focus on Spix’s disc-winged bat, *Thyroptera tricolor*, a small insectivorous species that roosts in the developing tubular leaves of plants in the order Zingiberales (Vonhof and Fenton 2004) in groups of approximately 5 individuals (Vonhof et al. 2004; Sagot et al. 2018). This species is known to use a call-and-response contact calling system for maintaining very stable group composition (Chaverri 2010) despite moving among roost-sites on a daily basis. Spix’s disc-winged bats produce two different types of social calls: the “inquiry” calls that are emitted by flying individuals and “response” calls that are emitted by roosting individuals in response to inquiry calls to guide and attract their conspecifics to the roosts (Chaverri et al. 2010). In this species, the rates of response call production are relatively consistent within, but vary widely among individuals (Chaverri and Gillam 2015). Furthermore, social groups are composed by a combination of vocal and non-vocal bats in the context of response calling, and thus around 50% of individuals produce response calls upon hearing inquiry calls from group and non-group members, whereas the rest never vocalize (Chaverri and Gillam 2015; Sagot et al. 2018).

We simulate vocal exchanges in *T. tricolor* to gauge the energetic costs of response call production. If individuals actively respond to the inquiry calls of their conspecifics, we expect metabolic rates to increase significantly; specifically, oxygen consumption should increase when bats vocalize for longer periods of time, as studies in a number of taxa demonstrate that vocalizations increase energy expenditure (Ryan 1988; Oberweger and Goller 2001; Ophir et al. 2010). We also test whether resting metabolic rates (RMR), i.e. those that reflect the metabolic rate of an individual during its inactive period (McNab 1997), correlate with response call production. Previous studies suggest that levels of activity or aggressiveness, which are traits that allow us to distinguish among animal personalities, are either positively or negatively influenced by resting metabolic rates (Careau et al. 2008). In the first case, termed the “performance model”, animals with greater levels of activity or aggression require larger organs to sustain these traits, and thus have higher-than-average maintenance costs (Daan et al. 1990). In contrast, the “allocation model” predicts a negative relationship between RMR and activity or aggressiveness because when food is limited, only individuals with lower self-maintenance costs can afford to spend energy in additional activities (Careau et al. 2008). While we have no a priori expectation regarding which model, performance or allocation, may predict response calling rates in *T. tricolor*, we test this to increase our understanding of the factors that may explain vocal personalities in the context of social communication.

## Methods

We collected data on metabolic rates for 38 individuals (18 adult females, 10 adult males, 3 subadult females, 4 subadult males and 3 juvenile males) from 11 social groups (i.e., individuals using the same roost at the same time) at Barú Biological Station in Southwestern Costa Rica, in July 2017. To find groups, we searched *Heliconia* spp., *Calathea* spp. and *Musa* spp. furled leaves, commonly used by *T. tricolor* as roosting sites (Vonhof and Fenton 2004). Once we located a roost, we captured all group members and placed them inside a cloth holding bag to bring them to the laboratory. Back in the laboratory, we weighted all the individuals and measured their forearm lengths (as a measure of body length). We also sexed, aged, and determined the reproductive condition for all bats captured.

For each individual, we were interested in two parameters: 1) Resting Metabolic Rate (RMR), and 2) metabolic rate while producing response calls. The animals were placed singly inside a tubular structure made of transparent plastic; there they remained safely attached to the interior’s smooth surface. The tube and bat were then placed inside a metabolic chamber and let to acclimate for 30 min. We measured the bats’ oxygen consumption using the methods described below, resting and while listening/responding to conspecific inquiry calls. All measurements were made in a silent room at ambient humidity (70%) and temperature (27°C) during daytime hours. At the end of the experiments, we provided mealworms (*Tenebrio molitor*) and water *ad libitum* to all individuals before releasing them in the same area where they were originally captured.

*Thyroptera tricolor* bats only produce response calls after an inquiry call has been emitted, and do so primarily during the day (Chaverri et al. 2010); thus, we broadcasted previously recorded inquiry calls to elicit response calling from the bats within the chamber. These inquiry calls were previously collected from five individuals flying within a large flight cage (3 x 4 x 9 m) for a total of 1 minute; none of these individuals were later included in our respirometry experiments. A total of 67 inquiry calls were identified in the 1-min recording, and the playback was continuously run for 10 minutes through an UltraSoundGate Player to a broadband loudspeaker (Ultrasonic Omnidirectional Dynamic Speaker Vifa, Avisoft Bioacoustics) placed inside the chamber. We recorded response calls produced by the individuals inside the chamber with an Avisoft condenser microphone (CM16, Avisoft Bioacoustics, Berlin, Germany) through Avisoft’s UltraSoundGate 116Hm onto a laptop computer running Avisoft-Recorder software (sampling rate 384 kHz, 16-bit resolution), placed also inside the chamber. We also video-recorded each of the trials to estimate the effect of movement (i.e. how long the bats were actively moving during the trials) for better interpretation of the metabolic rate results.

### Metabolic rate measurements

We measured O_2_ consumption (VO_2_) of each individual using an intermittent-flow-through respirometry. This set-up consisted of short-term trials (10 min) of closed respirometry followed by a flushing interval of 10 min that allowed the saturated air to be pumped out of the chamber and replaced by new air, avoiding CO_2_ accumulation. This method was used instead of a flow-through respirometry since it was not possible to measure flow rate. We placed each bat into a 2L acrylic chamber lined with paper to reduce sound disturbance (i.e., reduction of echo interference from playback). Air was pumped into the chamber using a standard fish tank pump and then drawn out and passed through a column of indicating Drierite TM connected to the ML206 gas analyzer fed from a damped, micro-vacuum pump (200 mL/min; ADInstruments, Bella Vista, NSW, Australia). Since we did not dry the air going into the chamber, we measured relative humidity of incurrent air with an electronic hygrometer, and mathematically scrubbed water vapor to provide a VO_2_ corrected to standard temperature pressure dry (STPD). We recorded the voltage outputs of the gas analyzer and thermocouple at a sampling frequency of 10 Hz using a PowerLab ML750 A/D converter (ADInstruments) and LabChart software (ADInstruments). For each bat we recorded O_2_ consumption for 10-min intervals of closed respirometry with and without sound broadcast. We calculated the whole individual metabolic rate (O_2_ ml h^-1^) using equation (4.9) of Lighton (2008), correcting for ambient pressure and standard temperature afterwards.

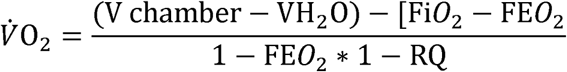

where V chamber is the volume of the chamber calculated by subtracting an approximation of the volume of the bat (mass multiply by 0.98) to the actual volume of the chamber (2L), VH_2_O is the water vapor in the chamber; Fi0_2_and FE0_2_ are the fractional concentration of O_2_ at the start and end of the experiment respectively. RQ is the respiratory quotient.

We converted oxygen consumption rate V O_2_ into energy expenditure in kJ by utilizing the oxy-joules equivalents (MR_kj_ in kJ hr^-1^) according to the following equation from Lighton (2008):

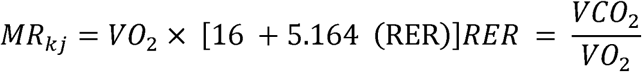

where RER is the respiratory exchange ratio (VCO_2_/VO_2_). We assumed a RER of 0.77, previously reported for insectivorous bats (Speakman et al. 1989b).

All sampling protocols followed guidelines approved by the American Society of Mammalogists for capture, handling and care of mammals (Sikes 2016) and the ASAB/ABS Guidelines for the use of animals in research. This study was conducted in accordance with the ethical standards for animal welfare of the Costa Rican Ministry of Environment and Energy, Sistema Nacional de Áreas de Conservación, permit no. SINAC-ACOPAC-RES-INV-008-2017.

### Data Analyses

We compared metabolic variables (i.e., RMR and energy expenditure during trials with sound) among age categories using a one-factor ANOVA and Tukey comparisons at an alpha level of 0.10. We found significant differences in RMRs between juveniles and adults, but not between adults and subadults (F_2,35_=2.95, P = 0.01). Therefore, we merged data for the latter but eliminated juveniles from further analyses. Our sample size for subsequent tests was 21 females and 14 males.

To determine if males and females differed in the amount of time spent producing response calls or moving, we conducted two separate Mann-Whitney U-tests, as the data were non-normally distributed. We also ran a Chi-square test to determine if the proportion of vocal (i.e., an individual that produced at least one response call) vs. non-vocal bats differed between males and females. We then determined if males and females differed in resting metabolic rate and metabolic rate while producing response calls with two separate independent samples t-tests. We analyze data separately for males and females as previous studies have shown that the strength and direction of selection on resting metabolic rates may differ according to sex (Burton et al. 2011).

To test if more vocal bats (i.e., bats that vocalized for longer periods of time) had higher metabolic rates, we conducted a linear model with energy expenditure in kilojoules (kJ) as the response variable, sex as a fixed factor, and as regressors, we selected the time the bats spent 1) moving, 2) producing response calls, 3) or other types of calls (echolocation, distress, and other calls of unknown function). We also included 4) mass as an additional regressor in the model. We retained the strongest explanatory variables using backward elimination and ran analyses separately for males and females. We also determined which was the variable with the greatest explanatory value based on the CP Mallows. We generated Q-Q and predicted vs. residual plots to test for normality and homogeneity of variances, respectively; both tests show that all assumptions of the model were met. In our results, we include the estimates of the multiple regression and those of simple regressions for explanatory variables kept in the model, to verify if independent effects on O_2_ consumption are positive or negative.

To determine if RMR is related with the time bats spend producing response calls, we conducted a generalized linear model with time spent producing response calls as the response variable, and energy expenditure (kJ) and sex (and their interaction) as fixed factors. The dependent variable was non-normally distributed (Shapiro-Wilks tests = all p-values < 0.001) and could be modeled best by a negative binomial distribution (p-value = 0.17).

Finally, we tested if sex and the propensity to produce response calls or not had an effect on the difference in energy expenditure during resting trials and during trials with sound through a general linear model. The difference in energy expenditure was estimated as the amount of kJ consumed during trials with sound minus the amount of kJ consumed during trials without sound. Bats were categorized as being vocal if they produced at least one response call during our trials with sound.

## Results

Bats were non-vocal during the 10-minute trials in which we measured the resting metabolic rates, i.e., those for which no sounds were broadcast. However, for trials in which we broadcast inquiry calls, bats vocalized for an average of 27.47 seconds (SD = 37.17); many individuals (n = 11) were non-vocal while the rest produced various types of vocalizations for up to 125.42 s. Animals produced three distinct calls with known functions and in decreasing order of frequency: response calls, which accounted for 61% of the time spent vocalizing; echolocation calls, which accounted for 21%; and distress calls (10% of the time). In some occasions, bats produced other calls with unknown function, which accounted for 8% of the time spent vocalizing (Fig. 1). There was no difference in the time spent vocalizing between males and females for any of the call types analyzed (P > 0.13), nor was there a difference in the proportion of vocal vs. non-vocal individuals between males and females (vocal females = 38%, vocal males = 43%; P = 0.77). Time spent moving was also not significantly different between males and females (P = 0.72).

**Fig. 1.**
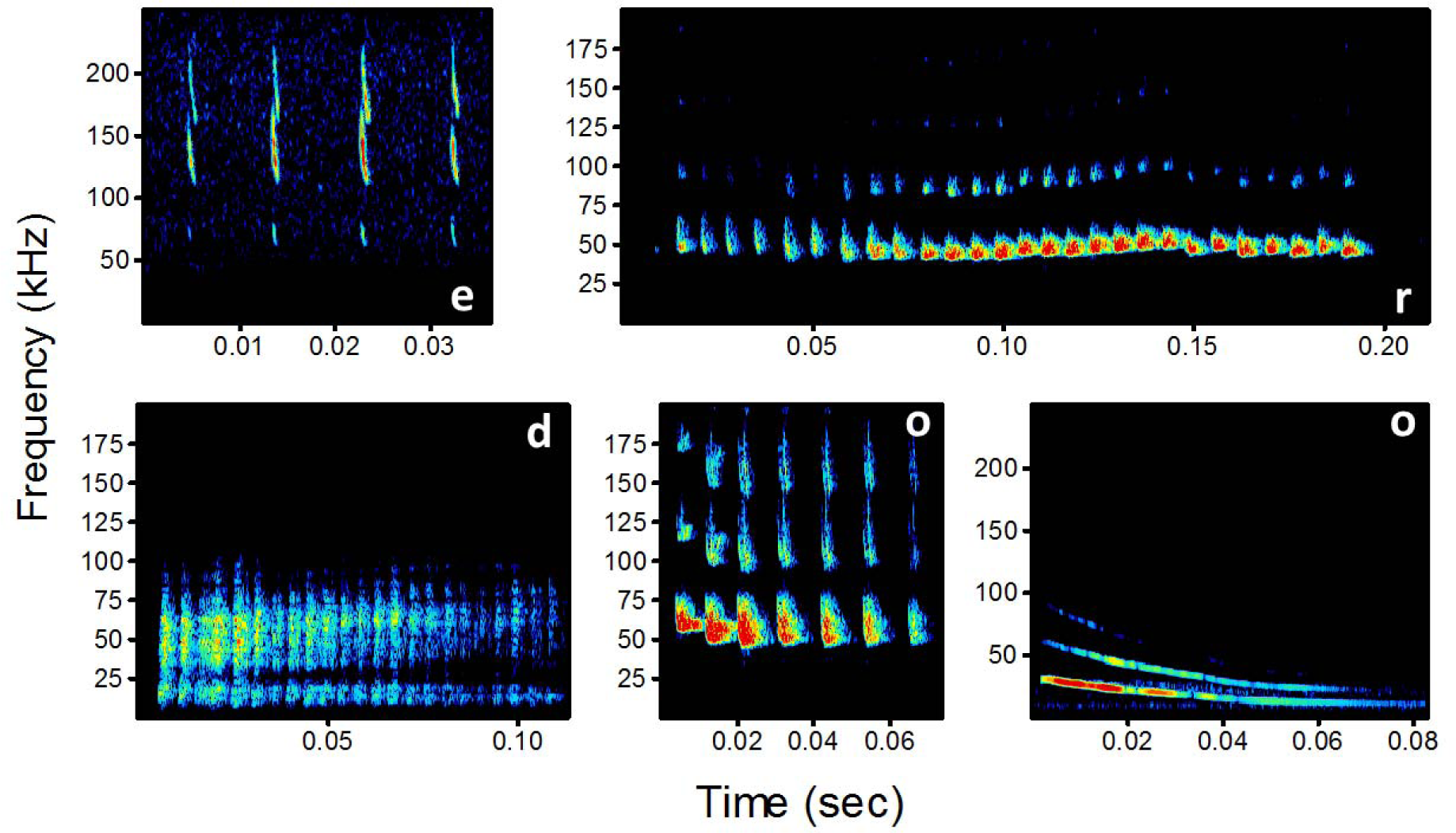
Sonograms depicting exemplars of call types recorded during our 10-min respirometry sessions: echolocation (e), response (r), distress (d), other (o).

Animals consumed an average of 7.80 ml O_2_ h^-1^ during trials when no sounds were emitted (Table 1). Females had a significantly greater energy expenditure (kJ) during periods of inactivity than males (t = 2.57, p = 0.01). During the experiments with sound, bats consumed an average of 16.93 ml O_2_ h^-1^. There was not a significant difference in energy expenditure (kJ) between males and females during trials with sound (t = 1.06, p = 0.29).

**Table 1.** Whole Animal Metabolic Rate (ml O_2_ h^-1^) during trials when inquiry calls were broadcasted (sound) or when bats were resting (no sound).

| Sex | Weight (g) | Sound |  | No sound |  |
| --- | --- | --- | --- | --- | --- |
| | | Range | Mean±SD | Range | Mean±SD |
| Female | $4.50 \pm 0.40$ | 8.37-38.89 | $17.94 \pm 7.40$ | 3.38-16.29 | $8.86 \pm 3.62$ |
| Male | $4.10 \pm 0.36$ | 4.80-30.13 | $15.42 \pm 6.74$ | 2.30-10.95 | $6.22 \pm 2.33$ |
| All | $4.32 \pm 0.43$ | 4.80-38.89 | $16.93 \pm 7.15$ | 2.30-16.29 | $7.80 \pm 3.39$ |

The results of our general linear model, where we tested if sex and being vocal had an effect on the difference in energy expenditure during trials with sound compared to resting trials, show that the latter factor (being vocal) had a significant effect (F_1,31_ = 11.70, p < 0.01). The average increase in energy expenditure for vocal bats during trials with sound was 12.42 kJ (±1.48), whereas the increase for silent bats was 4.48 (±1.78; Fig. 2). Although the difference in energy expenditure for vocal vs. non-vocal individuals was greater for males than for females, neither sex nor the interaction between sex and vocal behavior was significant (p > 0.25).

**Fig. 2.**
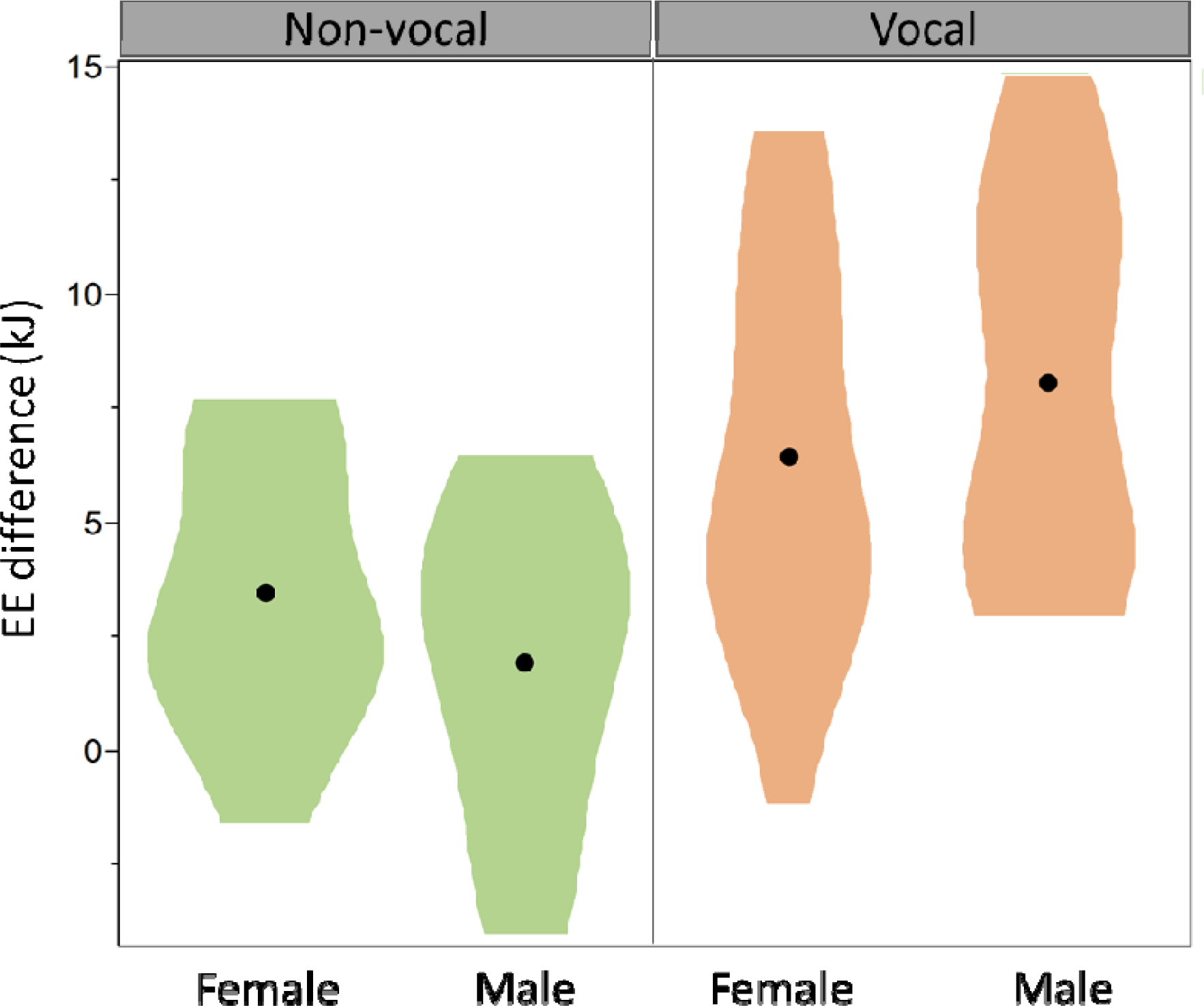
Difference in energy expenditure, measured as the difference in kJ during trials with sound minus kJ during trials without sound, for vocal and non-vocal males and females.

For females, our multiple linear regression model indicates that energy expenditure (kJ) was significantly and positively influenced by the time spent producing calls, such as echolocation and distress; however, time spent producing response calls did not affect their energy expenditure, nor did time spent moving and body mass (Table 2, Fig. 3). In the model for males, there was a significant and also positive effect of time spent producing response calls on energy expenditure; thus, males that produced more response calls had greater energy expenditure. Time spent moving also contributed to an increase in energy expenditure in males (Table 2, Fig. 3).

**Table 2.**
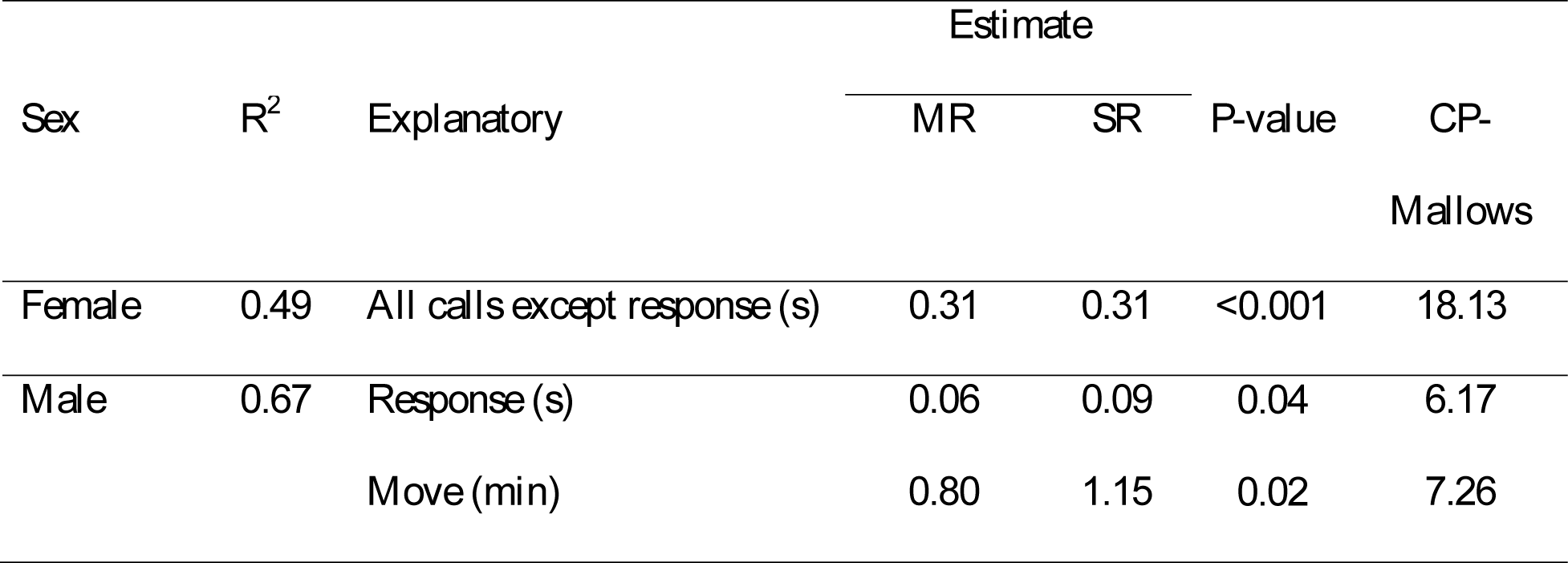
Results of the multiple regression analysis with energy expenditure (kJ) as the response variable and several explanatory variables. We include the estimates of the multiple regression (MR) as well as those of the simple linear regression (SR). The CP-Mallows indicates the explanatory power of the variables included in the model.

**Fig. 3.**
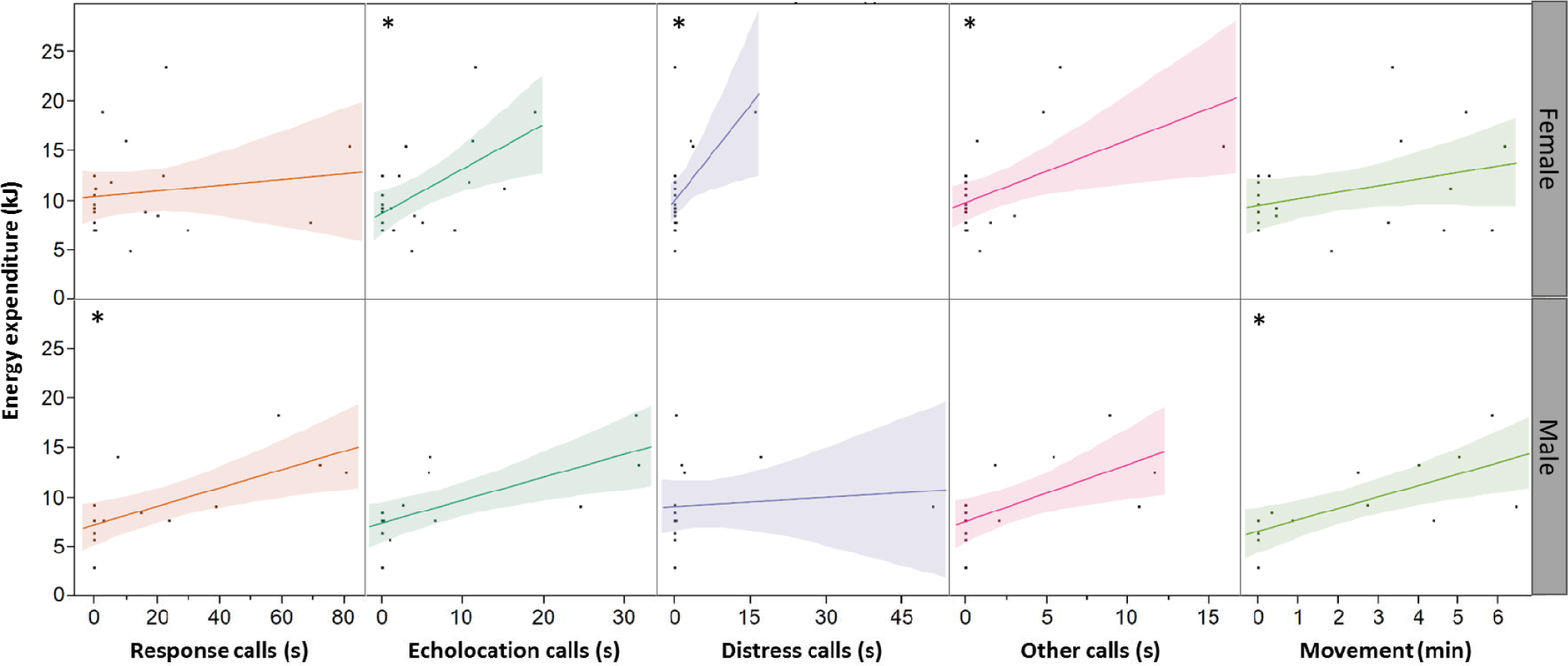
Scatter plots showing the relationship between time invested in various activities (emission of several types of calls and movement) and energy expenditure. Upper plots show the results for females and lower plots results for males. Asterisks indicate significant relationships according to the multiple regression analysis (see table 2).

Time spent producing response calls was significantly influenced by the interaction between sex and RMR (F_1,31_=5.05, p-value = 0.03), according to our generalized linear model. When performing the model separately for males and females, the relationship between RMR and time producing response calls for females was non-significant (p-value = 0.65), whereas for males the relationship was negative and significant (p-value = 0.003; Fig. 4). Thus, as RMR decreases in males, there is an increasing probability that they will emit response calls for longer periods of time.

**Fig. 4.**
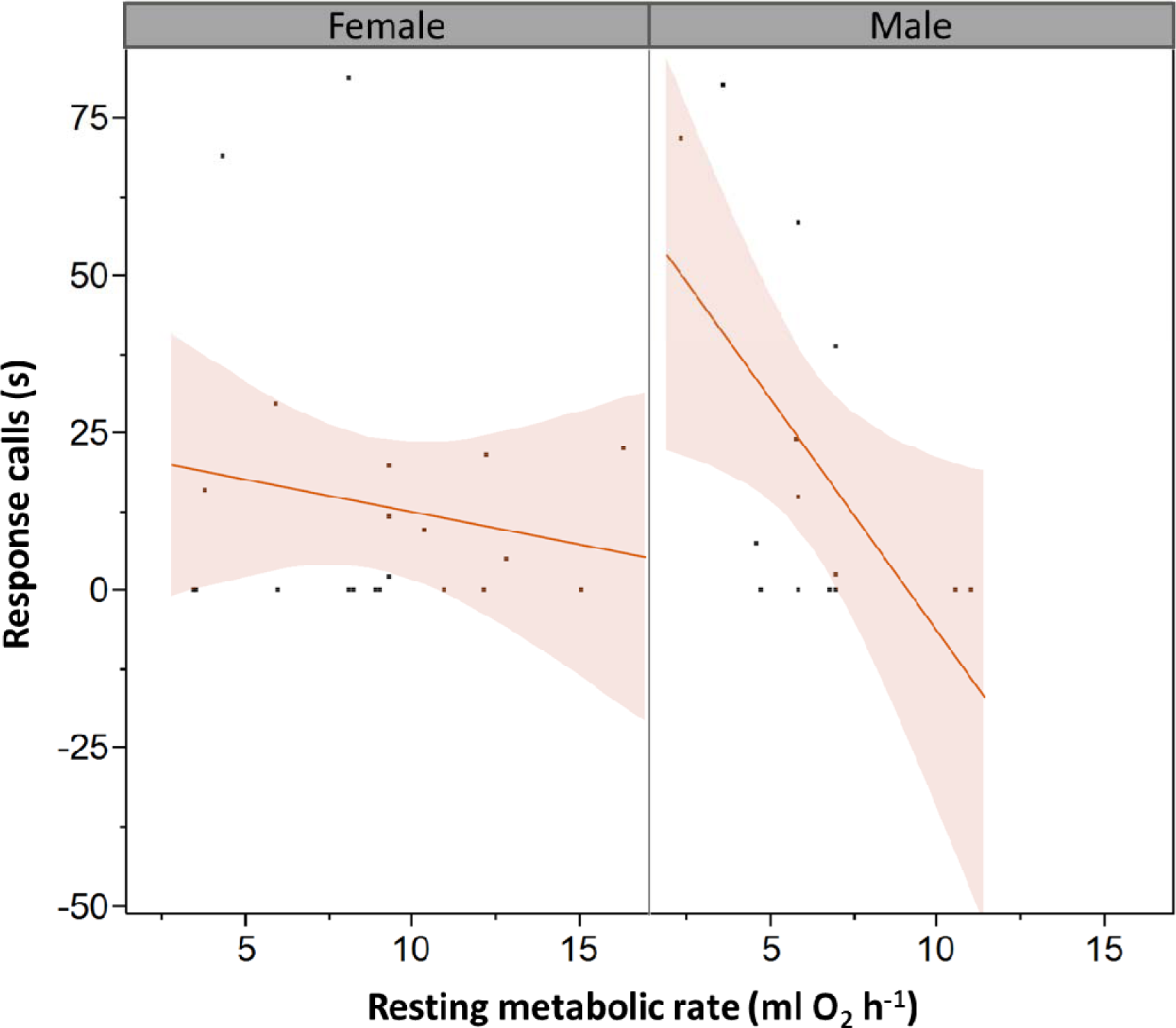
Relationship between RMR and the time spent producing response calls (in seconds) for males and females. The shaded area around the trendline shows the 95% confidence interval.

## DISCUSSION

Our results demonstrate that the production of social calls that are used to indicate the position of a roost site increases the energetic expenditure of bats. By producing even just one response call upon hearing an inquiry call, individuals significantly increased their resting metabolic rate. For males, the time spent producing response calls had a positive effect on energy expenditure, which did not occur in females, despite the fact that males and females produced a similar number of response calls and both were vocal in similar proportions. Vocal females increased their metabolic rate, on average, 1.4-fold when producing response calls, whereas males experienced a 3.2-fold increase.

Vocal communication can be observed in every major taxonomic group and in virtually every environment, and it is energetically demanding for many species (Ryan 1988; Prestwich 1994; Oberweger and Goller 2001). Birds, for instance, increase their metabolic rate at least 2.5-fold when producing courtship calls, while ectotherms such as insects and amphibians can exhibit an 8-fold increase (Ophir et al. 2010). This is because sound production elevates muscular activity (Prestwich 1994; Gillooly and Ophir 2010), and increases the vibration frequency of the muscles that produce the sounds, elevating metabolic rates (Skoglund 1961; Martin 1971; Elemans et al. 2004). In *T. tricolor*, both males and females significantly increased their metabolic rates while producing response calls, suggesting that energetically compromised bats cannot afford extra energy expenditures in functions that are not part of their normal daily maintenance activities. This might help us explain why many individuals are non-vocal (Chaverri and Gillam 2015; Sagot et al. 2018). Furthermore, in males but not in females, the increase in metabolic rate was proportional to the time spent vocalizing, suggesting that males that are energetically limited cannot produce response calls, or can only vocalize for short periods of time. This is possibly the reason why, in our study, only a small proportion of males produced vocalizations during relatively large amounts of time. This has also been found in other species such as bottlenose dolphins and birds, in which oxygen consumption increases with song duration and call rate (Oberweger and Goller 2001; Franz 2003; Noren et al. 2013).

Although sound production can be energetically demanding, in some species this activity does not increase an individual’s metabolic rate (Ilany et al. 2013). For example, male hyraxes (*Procavia capensis*) that sing more, counterintuitively conserve more energy. Likewise, echolocating bats do not significantly increase their energy expenditure during flight (Speakman et al. 1989a; Voigt and Lewanzik 2012). However, even when vocalizations are not energetically demanding, they can still be considered a handicap (Gil and Gahr 2002). This is because producing these signals requires time, learning and specialized structures, and it can increase the chances of being detected by prey and predators (Koren and Geffen 2009; Charlton et al. 2011; Wyman et al. 2012).

We also found that differences in RMRs may predict the time spent producing response calls in males. Specifically, we found that males with lower RMRs emit response calls during longer periods of time. These results confirm that levels of activity, in our case measured through the time spent vocalizing, are negatively influenced by RMR, which provides support for the allocation model. This model predicts that only individuals with lower self-maintenance costs can afford to invest part of their daily energy budget in additional activities (Careau et al. 2008). Despite our results, the most common trend in vertebrates is for RMR to positively influence activity, thus supporting the performance model; however, males often exhibit the opposite trend, which might indicate that they produce signals with enough energy to experience a trade-off between RMR and activity (Stoddard and Salazar 2011). This latter argument might explain the differences in energetic expenditure during response calling observed for males and females in our study. For instance, response calls in males might not only play a role in cooperative signaling of roost location (Chaverri and Gillam 2010), but may also function for mate attraction; if so, males could be under strong selection to produce high quality/energy calls as an honest signal of their body size and condition (Schuchmann and Siemers 2010). Thus, the physiological explanation for the differences in energetic costs of social signaling between males and females could be hormonal, as several studies demonstrate that male sexual hormones significantly alter the relationship between resting metabolic rates and signal quality or levels of activity (Wikelski et al. 1999; Lynn et al. 2000; Buchanan et al. 2001). Future studies should try to confirm the link between acoustic features of social calls like maximum energy, metabolic cost and mating success, in addition to addressing the potential role of response calls for mate attraction in *T. tricolor*.

In conclusion, our study demonstrates for the first time that social calls increase energetic expenditure in bats. Given that bats depend so strongly on acoustic signals for modulating multiple social activities (Gillam and Fenton 2016; Chaverri et al. 2018), our findings suggest that energetic trade-offs may be of particular importance to understanding communication in this group of mammals. The results of our study will surely extrapolate to various other species in diverse contexts; however, it is the differences among systems that seem most fascinating. In our case, we have addressed the costs of acoustic signaling during contact calling, but further studies could reveal interesting tradeoffs for signals such as those employed between mothers and offspring, or between males and females in the context of mate attraction, among others. Finally, our results provide a step forward in our understanding of how physiology modulates behavior. For example, many studies demonstrate that there is a link between resting metabolic rates and various personality traits (Careau et al. 2008; Careau and Garland 2015). Incorporating physiological trade-offs to studies of animal personalities in the context of communication may allow us to understand many aspects of social aggregations, including social roles and communication networks.

## Acknowledgements

The authors would like to thank Tenaja Smith-Butler and Cayla Turner for their help during field work and for video analyses, and Ronald Villalobos for logistics support. We also thank Julio Bustamante and Lilliana Rubí Jimenez for their help during research permit application. Finally, we thank the Centro Biológico Hacienda Barú for their continuous support of our research.

## Data availability

The data supporting this article are available from the Figshare Digital Repository: https://doi.org/10.6084/m9.figshare.13003805.v1.

